# Pigment Binding in The Light-Dependent Protochlorophyllide Oxidoreductase

**DOI:** 10.1101/2023.08.28.555081

**Authors:** Penelope Pesara, Katarzyna Szafran, Henry C. Nguyen, Abhishek Sirohiwal, Dimitrios A. Pantazis, Michal Gabruk

## Abstract

The Light-Dependent Protochlorophyllide Oxidoreductase (LPOR) is a key enzyme in chlorophyll biosynthesis and its photocatalytic mechanism has long intrigued researchers. However, the lack of structural data for the active complex has impeded understanding of its reaction mechanism. A recent high-resolution structure of enzyme in the active conformation has established a robust foundation for validating hypotheses concerning pigment binding, residue involvement, and consequently, the reaction mechanism. Surprisingly, this new structure challenges previously proposed mechanisms, especially concerning the orientation of the bound protochlorophyllide (Pchlide) pigment. In this study, we employ molecular dynamics and hybrid quantum-mechanics/molecular-mechanics (QM/MM) simulations along with site-directed mutagenesis to compare two Pchlide binding modes: one aligned with previous proposals (mode A), and the other consistent with the recent experimental data (mode B). Binding energy calculations reveal thermodynamic instability of binding mode A due to nonspecific interactions, while mode B exhibits distinct stabilizing interactions yielding favorable binding. QM/MM-based local energy decomposition analysis unravels a complex interaction network that reinforces pigment stabilization in this conformation. Notably, interactions involving Tyr177, His319, and the carboxyl group at C13^1^ influence the pigment’s excited state energy and potentially contributing to the substrate specificity of the enzyme. Our results uniformly favor binding mode B as represented in the new cryo-EM structure, over the previously assumed mode A. These findings challenge established interpretations and underscore the need for a comprehensive re-evaluation of the reaction mechanism of LPOR that correctly considers pigment interactions and substrate orientation in the binding pocket.

**Significance Statement:** A crucial step in the biosynthesis of the all-important photosynthetic pigment chlorophyll is the reduction of a double C=C bond in its precursor protochlorophyllide (PChlide). This is catalyzed by the Light-Dependent Protochlorophyllide Oxidoreductase (LPOR) via an extremely rare example of a biological photocatalytic reaction. Understanding of the LPOR mechanism has been hindered by limited insight into the structure of its active complex. A recent high-resolution LPOR cryo-EM structure substantiates pigment binding, residue interactions, and the reaction mechanism, but contrasts markedly with all previous assumptions regarding the binding mode of the substrate PChlide. Using molecular dynamics simulations, quantum-mechanics/molecular-mechanics calculations, and mutagenesis, we compare and evaluate the two possible Pchlide binding modes, the one assumed previously (mode A) and the one supported by recent data (mode B). Our findings conclusively favor mode B, challenging prior assumptions and pointing toward novel mechanistic possibilities for this unique photocatalytic reaction.

## Introduction

Chlorophyll (Chl) is the most essential natural pigment in organisms that carry out photosynthesis, the process of capturing sunlight and transforming it to chemical energy that powers Earth’s biosphere (1). A crucial step in chlorophyll biosynthesis is the light-dependent conversion of protochlorophyllide (Pchlide) to chlorophyllide (Chlide) by reduction of the C17=C18 double bond of the former (2). This reaction, a rare example of light-driven biosynthetic transformation, is catalyzed by the light-dependent protochlorophyllide oxidoreductase (LPOR) that employs NADPH as a cofactor. Owing to the light-driven nature of this reduction, Pchlide has also a regulatory role in Chl biosynthesis (3, 4). The mechanism of action of LPOR remains poorly understood (2, 5, 6), because of challenges related to the light-activated nature of the reaction, the involvement of several experimentally ill-characterized intermediates, and the lack of accurate structural information regarding the active form of the enzyme and the precise positioning of the substrate and cofactors.

An early NMR study by Begley and Young suggested that the *pro*-S face of NADPH delivers a hydride to C17 of Pchlide; while an amino acid residue is assumed to subsequently protonate C18 to complete the reaction (7). A suitable candidate for this residue was proposed based on the sequence similarity between LPOR and short-chain dehydrogenase reductase (SDR) family (8). Most enzymes in this family share a conservative catalytic motif that consists of a key Tyr276 and Lys280 residues (the numeration for PORB isoform of *A. thaliana*)(9). The classical SDR mechanism employs the interaction between Tyr and Lys residues to lower the tyrosine hydroxyl pKa so it can serve as proton donor. Numerous site-directed mutagenesis studies confirmed the importance of Tyr276 for the activity of LPOR (10, 11). Importantly, in all published experiments the mutation of Tyr276 only *lowered* and did not abolish LPOR activity as would be expected for the above assumed role. Nevertheless, the accumulated experimental data allowed the proposal of a plausible picture of the active site of the enzyme (mode A, Fig. 1B), which served as a starting point for several attempts at computational modeling of mechanistic scenarios. In these studies, either a homologous model of the enzyme or some experimentally derived structures were employed (6, 12, 13). Crucially, however, in all these studies the conformation of the enzyme in the hypothetical active state and the binding mode of the substrate, Pchlide, were obtained with molecular docking techniques or were presumed to accommodate past assumptions regarding the stereospecificity of the reaction, the involvement of Tyr276, and the orientation of NADPH. Such binding mode was also assumed in two recent computational studies on the mechanism of LPOR, although the actual conclusions in these studies diverge drastically on the nature and sequence of events (6, 13).

In 2021 the first atomic structure revealing the architecture of the active site of the protein with bound Pchlide was reported (14). This reconstruction revealed that Pchlide is embedded within a helical lattice partially in the outer leaflet of the membrane sandwiched between the helix α10 and a region of the LPOR-specific loop (“the Pchlide loop”). The resolution of the protein within the structure allowed to determine the positions of all amino acid residues, including these responsible for Pchlide binding. Intriguingly, the conformation of the protein in the active state is nearly identical to the other LPOR structures obtained for the protein lacking the pigment, *except* for parts interacting with Pchlide. The two parts of the enzyme sandwiching Pchlide show two distinct conformations depending on the presence of the pigment. These conformations may represent two states of the enzyme: the *apo* and the *holo* forms, of which the latter is active. Crucially, even though the resolution of the electron density found in the pigment binding pocket of the active form of the enzyme was not sufficient to propose the orientation of Pchlide with complete confidence (14), the best fit of the pigment to the electron density clearly disagrees with the conventional assumptions on the mechanism of the reaction (Fig. 1A–C). Specifically, the nicotinamide ring of NAPDH and the Tyr276 are found in neither the proximity nor the orientation previously assumed with respect to C17 and C18, respectively (mode B, Fig. 1C). This, in turn, suggests a different mechanistic pathway than what has been discussed in the past.

The two distinct binding modes of Pchlide necessarily lead to distinct mechanistic scenarios. In the case of binding mode A, two mechanisms were proposed by Johannissen et al. (6) and by Silva and Cheng (13). Both suggest a light-activated reaction where Pchlide absorbs a photon that triggers a different reaction cascade. Johannissen et al. proposed a hydride transfer (HYT) from NADPH to C17 of Pchlide followed by proton transfer (PT) from a tyrosine residue to C18 (6). In contrast, Silva and Cheng supposed an initial electron transfer (ET) followed by PT from a tyrosine residue to C18, after which the HYT from NADPH to C17 would take place (13). The involvement of a cysteine residue was additionally introduced. The new cryo-EM structure, however, raises questions about the validity of all these models and corresponding mechanisms, owing to the fundamentally distinct positioning of Pchlide (mode B), which even challenges the conventional interpretation of the Begley–Young experiment (7), of a mechanism where NADPH is supposed to reduce C17. Nguyen et al. therefore hypothesized other alternative proton donors or water molecules, which cannot be seen using cryo-EM techniques, involved within the reaction mechanism in such a way that C18 can be reduced by cofactor NADPH.

The question on the mechanism of action of LPOR remains wide open. Given the central importance of the binding mode of Pchlide in addressing it, here we evaluate the validity of both competing modes A and B taking advantage of the new structure of the enzyme that depicts LPOR in its active form and combining a range of new computational and experimental data.

## Results

### Computational Analysis of Protochlorophyllide Binding

To investigate the two potential binding modes A and B of Pchlide within the substrate pocket of LPOR, we conducted a comprehensive analysis using two distinct computational methodologies encompassing classical atomistic molecular dynamics simulations and hybrid quantum mechanics/molecular mechanics (QM/MM). Our aim is to determine the Pchlide binding affinity in both modes and further gain atomic-level insights into the functional role of the protein in substrate stabilization. The first approach involved a comprehensive MM-PBSA (Molecular Mechanics Poisson–Boltzmann Surface Area) based characterization, enabling the computation of the binding energy along the dynamic progression while explicitly considering protein flexibility. In case of the MM-PBSA approach, the binding energies were averaged over a large ensemble of snapshots (see Supporting Information). The second approach employed a QM/MM technique combining the domain-based local pair natural orbital (DLPNO) variant of the coupled cluster method with singles, doubles and perturbatively included triples excitations (CCSD(T)) (15–19) as the basis for the Local Energy Decomposition (LED) scheme (15, 20, 21), which enables decomposition of the DLPNO-CCSD(T) interaction energy into fragment-pairwise additive contributions. The latter approach assesses the interaction energies between substrate and the protein by explicitly considering the quantum interactions among electron orbitals localized on distinct fragments.

**Figure 1.**
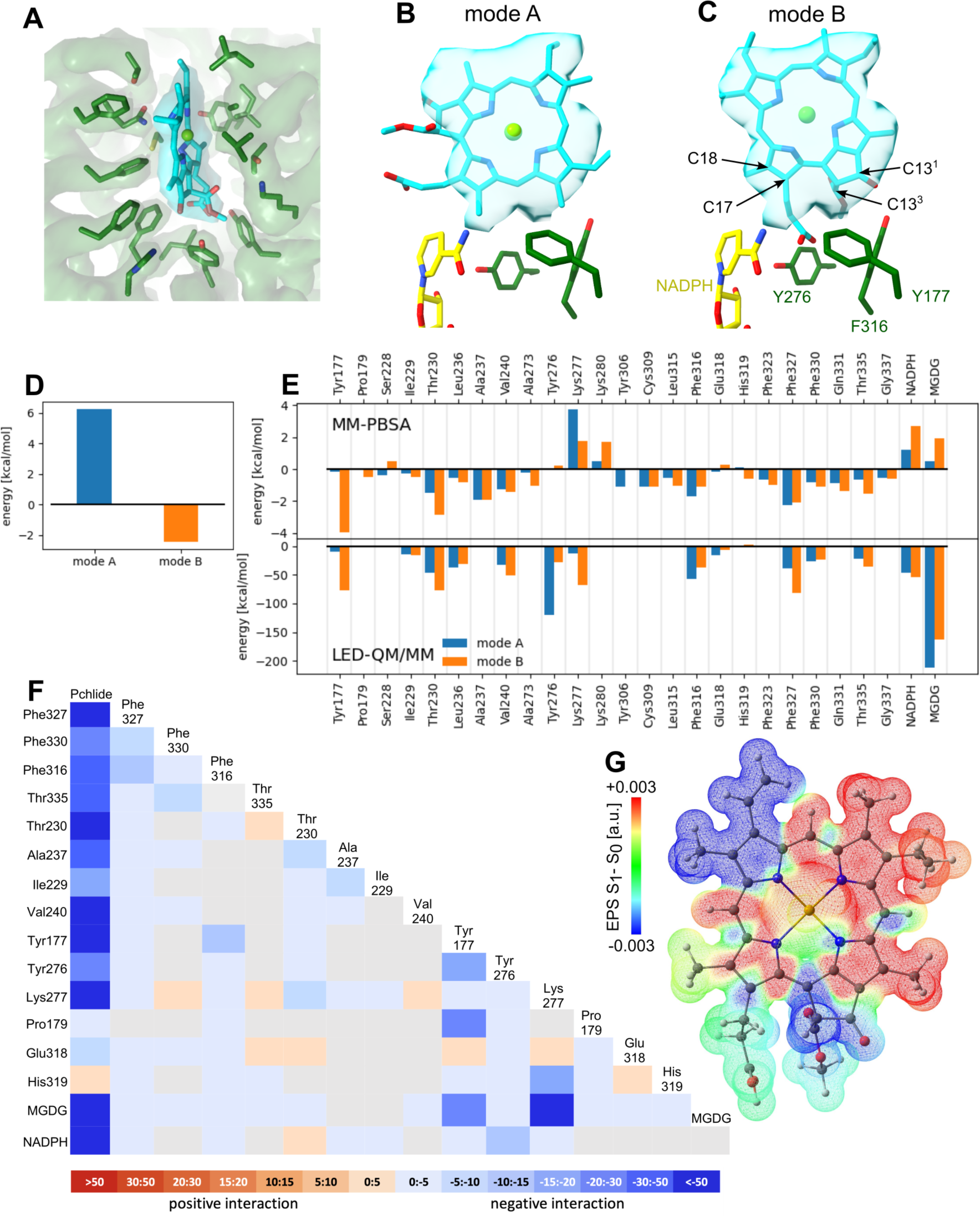
Two modes of Pchlide binding. **A.** Depiction of bound Pchlide (mode B) and corresponding cryo-EM density along with the molecular details of the residues in the binding pocket as resolved in the active form of LPOR (PDB ID: 7KJ9). Residues of Pchlide binding pocket with an electron density of the pigment. **B** and **C.** Pchlide binding modes A and B with the electron density of the pigment, respectively. **C.** Total Pchlide binding energy calculated with MM-PBSA approach. **D.** The contribution of selected residues to the Pchlide binding energy as calculated with MM-PBSA and LED QM/MM approaches. **E.** LED interaction map for mode B. **F.** Difference electrostatic potential map (S1 minus S0) associated with the S0→S1 (Q*y*) excitation for Pchlide in mode binding B, computed by QM/MM calculations using the ωB97X-V/def2-TZVP level of theory.

MM-PBSA binding energy computations clearly reveal stronger binding affinity for mode B of the Pchlide (Fig. 1D). The binding energy of Pchlide in mode A was found to be net positive, i.e., endergonic (+6.3 kcal/mol), whereas binding in mode B was exergonic (-2.4 kcal/mol) (Fig. 1D). To gain more molecular-level insights, the binding free energy was decomposed into per-residue contributions to identify which residues are important for the binding process within each mode. For mode A, the most significant contributors were Thr230, Ala237, Val240, Tyr306, Phe316, Phe327, Phe330, Gln331, and Thr335. For mode B, the key residues were Tyr177, Thr230, Ala237, Val240, Cys309, Leu315, Phe316, Phe323, Phe327, Phe330, Gln331, and Thr335 (Fig. 1E). Most of these residues exhibited stronger interactions for mode B, except for Phe316 and Phe327, where the interactions were predicted to be stronger for mode A. Overall, the amino acid residues in the substrate binding pocket significantly contribute to the overall binding energy (see Fig. 1) and exhibit differential stabilization between the two modes of Pchlide binding, particularly favoring mode B.

To go beyond the classical mechanics picture and further elucidate the energetics of Pchlide binding at the quantum chemical level, we employed a QM/MM-based LED analysis (Fig. 1E, SI appendix, Fig. S1). We found that for mode A the binding arises primarily from interaction terms with Tyr230, Tyr276, Phe316 and Phe327, as well as with MGDG (Fig. 1D, SI appendix, Fig. S1). In contrast, for mode B, LED analysis reveals a complex network of interactions that effectively stabilize the pigment in this specific conformation (Fig. 1E). On one side of the pigment we observed notable electrostatic and exchange interactions between the tetrapyrrole ring and three phenylalanine residues (316, 327, and 330) as well as Thr335 (Fig. 2A). On the other side of the pigment, a chain of hydrogen bonds involving Thr230, Lys277, and MGDG facilitate the connection between the two carbonyl oxygens of the Pchlide molecule at carbon atoms C13^1^ and C13^3^ (Fig. 2B). The LED analysis suggests that these interactions primarily arise from electrostatic and exchange forces, but electrostatic interactions dominate the MGDG–Pchlide interaction.

**Figure 2.**
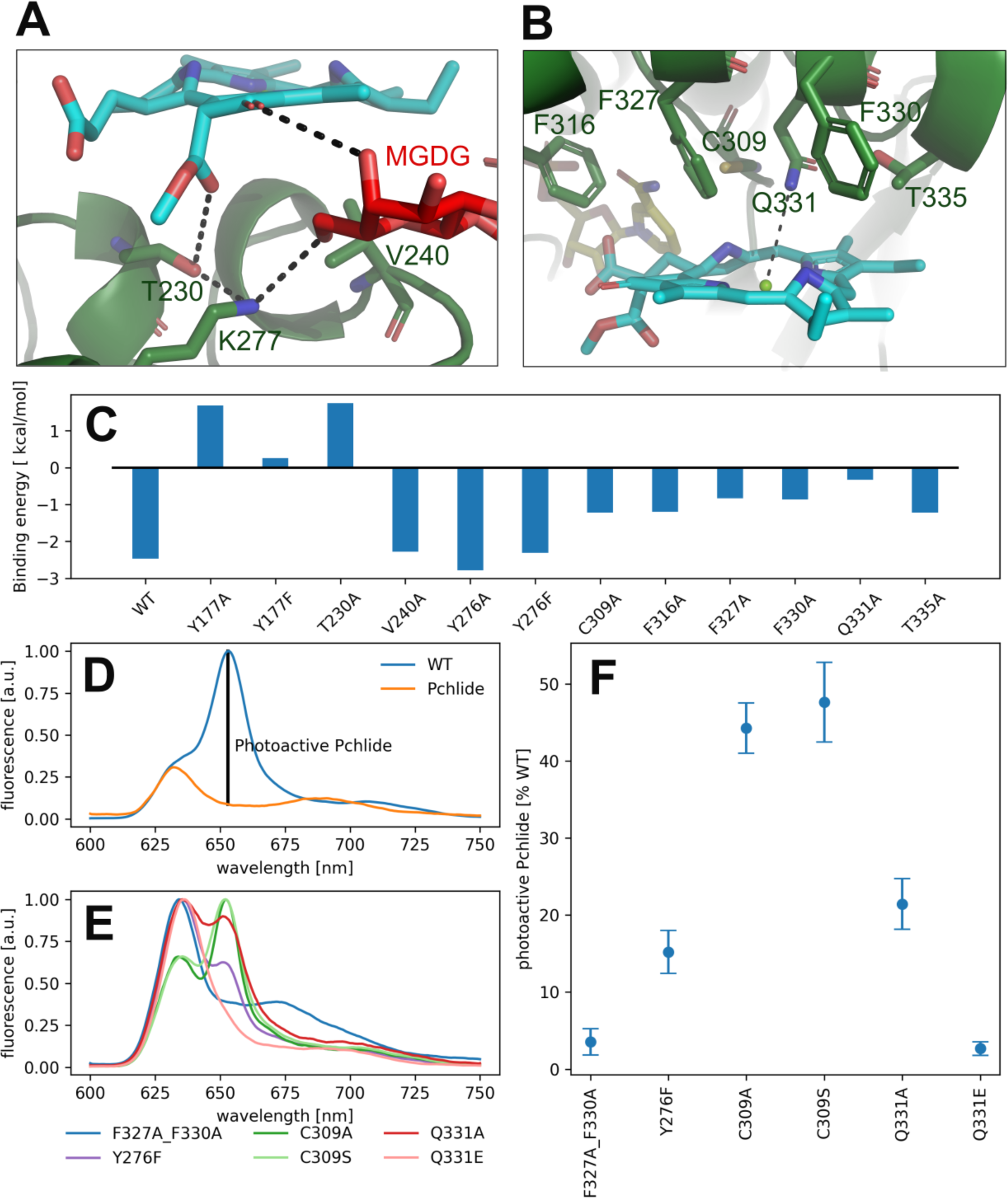
Pchlide binding in mode B. **A** and **B.** Major interactions between Pchlide, amino acid residues and MGDG. **C.** Impact of point mutations on binding energy assessed via the MM-PBSA methodology**. D** and **E.** The low-temperature fluorescence spectra of free Pchlide in a buffer (D: Pchlide), Pchlide in the reaction mixture with LPOR-WT, NADPH and lipids (D: WT), and Pchlide in the same conditions but with different mutants (E). The fluorescence intensity assigned to photoactive Pchlide in marked (D). **F.** The amount of photoactive Pchlide compared to WT-LPOR calculated for different mutants.

Polarization along the C17-C18 bond in the excited state is a key driving force for the proton-coupled electron transfer process. Therefore, we assessed the possible impact of the different Pchlide modes on this bond. Our excited state TD-DFT computations reveal that the polarization of the C17-C18 bond (judged by the S_1_-S_0_ ESP difference, Fig. 1G) shows similar behavior in mode A and mode B. In fact, gas-phase computations (see SI appendix, Fig S2) revealed similar bond polarization, implying that the excited-state polarization of this bond is intrinsic to the electronic structure of the Pchlide. Therefore, the different binding modes do not affect the intrinsic properties of the pigment and the fundamental features of light-driven excitation as judged by the S_1_–S_0_ ESP difference, but they do have implications on the nature of subsequent mechanistic events. Overall, and despite the fundamentally distinct theoretical foundations of the molecular mechanics and the correlated wave function approaches employed here, all theoretical approaches favor binding of Pchlide in the mode B consistent with the recent cryo-EM data rather than in the previously hypothesized mode A, and moreover identify a similar set of residues as contributing to the favorable binding in mode B.

### Site-directed mutagenesis and validation of the Pchlide binding mode

To further evaluate the influence of specific residues on the binding affinity of Pchlide in mode B, we conducted a computational (using MM-PSBA) and experimental mutational screening. We selected residues with significant contributions to the binding energy and performed selective substitutions (Fig. 2C) in order to examine the corresponding changes in the binding free energy (ΔΔ*G*). Among the chosen mutations, the system with Y177A, Y177F and T230A led to a destabilization of Pchlide binding, resulting in unfavorable binding energies (Fig. 2C). Conversely, the mutations of V240A, Y276F, Y276A exhibited minimal to negligible effects on the binding energies. Remarkably, the substitutions of C309A, Q331A, and the three phenylalanine residues: F316A, F327A, and F330A, led to a significant decrease in binding energies (Fig. 2C). Overall, these findings underscore the importance of specific residues in modulating the binding affinity of Pchlide in mode B and provide valuable insights into the molecular mechanisms governing this process.

To experimentally validate the computational predictions regarding the contributions of these residues to Pchlide binding, we performed site-directed mutagenesis on selected residues. Based on the available data and structural information, we chose to study the following mutants: Y177F, Y276F, C309A, C309S, Q331E, Q331A, and the double mutant F327A_F330A. To evaluate the impact of these mutations on Pchlide binding, we compared the low-temperature fluorescence emission spectra of Pchlide in a complex with lipids, NADPH and AtPORB: the WT and the respective mutants (Fig. 2D–F). The spectra of free Pchlide and the pigment bound to LPOR exhibit distinct differences, with maximum emission peaks separated by 23 nm (Fig. 2D). These spectral variations have been extensively employed to estimate the quantity of photoactive Pchlide (i.e., the pigment bound to the enzyme, Fig. 2D) in etiolated seedlings (22, 23). In our study, we employed the same approach to estimate the amount of bound Pchlide in the mutant variants (Fig. 2EF). Our findings revealed that the double mutant F327A_F330A and the Q331E mutant were barely able to bind the pigment, while the Y276F and Q331A mutants exhibited binding at approximately 20% compared to the wild type. Additionally, the C309A and C309S mutants showed binding at around 45% compared to the wild type.

Our calculations predicted that the interaction between Y177 and Pchlide is one of the strongest in mode B (Fig. 1C). LED analysis predominantly attributes this interaction to electrostatic and exchange energies, which aligns with the cryo-EM structure. The structure indicates a potential hydrogen bond between the hydroxyl group of tyrosine and a C13^1^ carboxyl of the pigment (Fig. 3A). This interaction is only possible in binding mode B because, in the flipped orientation (mode A), the carboxyl oxygen points towards the solvent while Tyr177 points towards ring A of Pchlide (Fig. 1A).

We further performed mutation studies on Tyr177 to assess its role in the pigment binding. The Y177F mutant showed the ability to bind Pchlide and form complexes with NADPH and lipids, all of which exhibited enzymatic activity in the presence of NADPH and Pchlide (Fig. 3B, SI appendix, Fig. S3). Interestingly, the fluorescence emission maxima of Pchlide bound by the Y177F mutant were blue-shifted compared to the wild-type enzyme (Fig. 3BC). To explore further, we also mutated the adjacent residue His319 (Fig. 3A), even though it was predicted to interact weakly with Pchlide (Fig. 1D). The H319A mutant was also enzymatically active in the presence of NADHP and Pchlide, and capable of complex formation with NADPH and lipids (Fig. 3B, SI appendix, Fig. S2). Both mutations, Y177F and H319A, influenced the emission maxima of Pchlide in the complexes with LPOR, LPOR and NADPH, as well as LPOR:NADPH:lipids, but the strength of the observed effects were distinct for each mutant.

In the case of H319A, the emission maximum of the LPOR:Pchlide complex was blue-shifted compared to free Pchlide, while for Y177F, it was red-shifted, albeit not to the extent of the WT enzyme (Fig. 3BD). The addition of NADPH to the reaction mixture caused a red-shift of the maxima for both mutants, but it was less pronounced compared to the WT enzyme, with Y177F showing a milder effect than H319A (Fig. 3BD). As for LPOR WT, the addition of lipids containing MGDG to the reaction mixture, along with NADPH and Pchlide, further red-shifted the emission maximum of the pigment to the characteristic value of 655 nm and led to the formation of filamentous oligomers (Fig. 3D). The addition of the lipids to the reaction mixtures of the mutants also resulted in the further red-shift of emission but not as profound as for WT enzyme. The observed red-shift was less profound for Y177F and for H319A (Fig. 3BD).

Finally, we also characterized a double mutant, Y177F_H319A, which was found to be enzymatically active in the presence of NADPH and Pchlide, and capable of forming the complexes with Pchlide, NADPH, and lipids (Fig. 3B, SI appendix, Fig. S2). In terms of emission maxima of the complexes, the double mutant behaved similarly to the Y177F mutant, but with even less pronounced red-shifts, approximately by 2–3 nm compared to Y177F (Fig. 3C). We confirmed that the Y177F_H319A double mutant could form filamentous oligomers in the presence of Pchlide, NADPH, and lipids (Fig. 3D), although they were less organized than those formed by the WT and were not as abundant on the grid.

**Figure 3.**
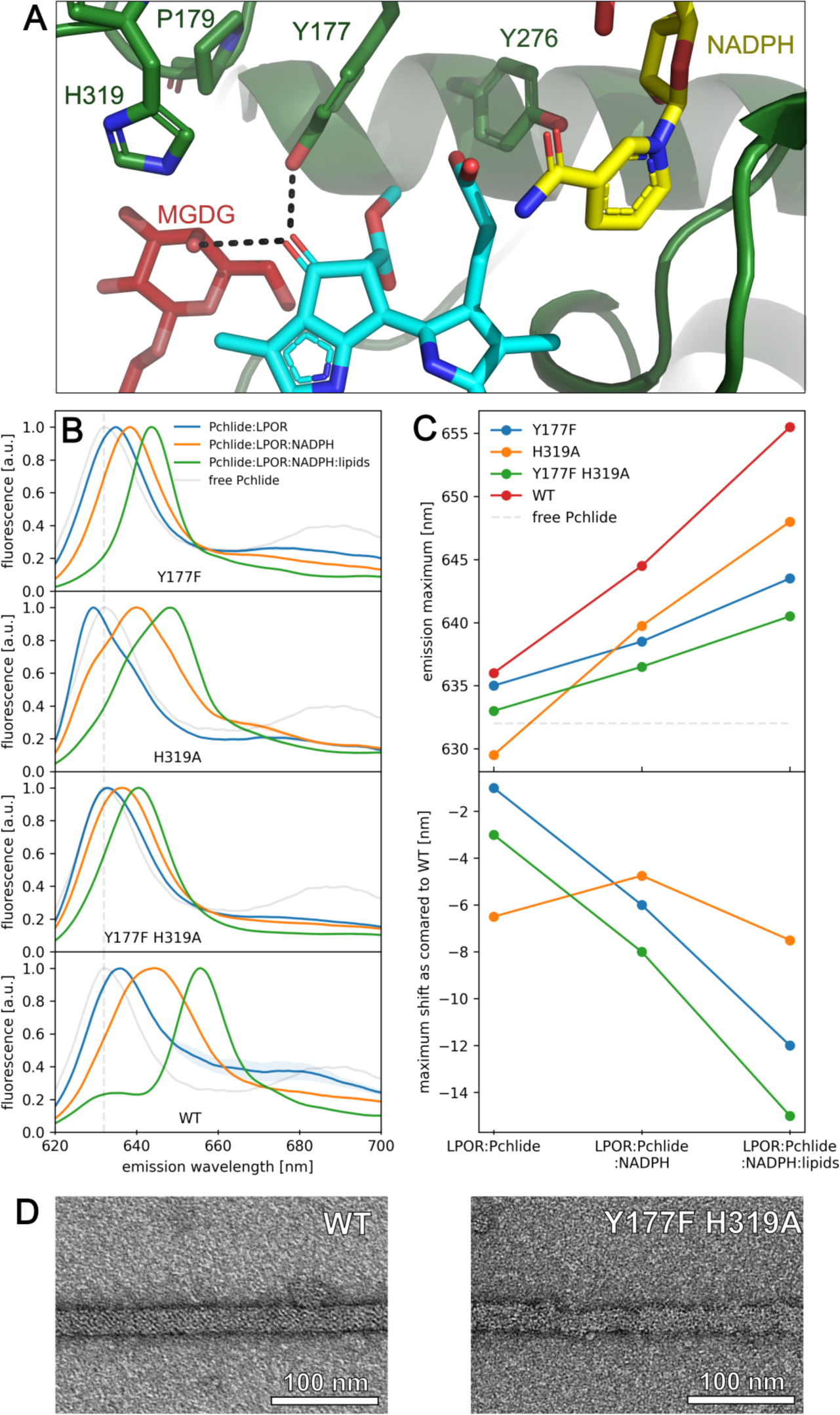
Hydrogen-bonding interactions with the keto group at C13^1^ of Pchlide affect the emission maximum of the pigment. **A.** Residues in proximity to keto group at C13^1^. **B.** Fluorescence emission spectra of free Pchlide in buffer and in complexes with LPOR WT and mutants. Reagents concentrations: 15 μM LPOR, 5 μM Pchlide, 200 μM NADPH and 100 μM lipids (50mol% MGDG, 35mol% DGDG, 15mol% PG). Spectra represent averages from at least two independent experimental replicates, with standard deviations depicted as bands. **C.** Fluorescence emission maxima of Pchlide in different complexes with WT LPOR and mutants and shifts of the emission maxima relative to WT enzyme. **D.** Negative-stain electron microscopy micrographs of LPOR:Pchlide:NADPH:lipids oligomers of WT enzyme and Y177F_H319A mutant.

## Discussion

The light-dependent catalysis carried out by LPOR has been a subject of great interest for many years. However, due to the limited availability of data on the structure of the active complex, investigations into the reaction mechanism have been built on uncertain foundations. Fortunately, with the recent determination of a high-resolution structure of the enzyme in its active conformation (14), we are now able to validate various hypotheses regarding pigment binding, the involvement of specific residues in this process, and ultimately, the reaction mechanism. This breakthrough provides a solid basis for advancing our understanding of the intricate workings of LPOR and its role in light-dependent catalysis.

In this paper we focused our efforts to compare two binding modes of Pchlide. We employ classical molecular dynamics and hybrid quantum-mechanics/molecular-mechanics (QM/MM) simulations to produce the “best-case” structural solutions for the two binding modes and we evaluate them using a range of criteria, including binding free energies determined using the MM-PBSA (24), and combine such simulations with experimental mutation data.

The MM-PBSA calculations performed for mode A indicate that the binding interaction lacks thermodynamic stability. A limited number of residues contribute to the binding energy; however, the observed interactions are nonspecific in nature. Previous simulations investigating this binding mode have also failed to identify any specific interactions with the pigment, except for interactions with the propionate group. These interactions were attributed to either Ser228 (Thr145 in T. elongatus LPOR) (25) or Ile229 (Val142 in Synechocystis LPOR) (13). In one of these studies, which provided relevant data, the remaining essential elements of the pigment were observed to either interact with water molecules (the magnesium ion, the C13^1^ keto group) or demonstrate a lack of interactions entirely (the C13^3^ keto group) (25).

Notably, any modifications to these pigment components have been shown to significantly impair or eliminate the affinity between modified Pchlide derivatives and LPOR, highlighting the enzyme’s sensitivity towards these parts of the pigment (25, 26). The enzyme’s exceptional substrate specificity is expected, considering that Pchlide is among several chlorophyll intermediates present in plastids. To uphold such high substrate specificity, LPOR must possess specific residues that interact exclusively with Pchlide, enabling the enzyme to discriminate it from other similar intermediates.

In binding mode B, we have discovered a distinct ensemble of interactions between the residues of LPOR and the pigment. Collectively, these interactions result in a negative binding energy, indicating thermodynamic stability. Notably, a subset of these interactions involves the vital components of Pchlide, namely the keto groups at C13^1^ and C13^3^, suggesting their potential contribution to the substrate specificity of LPOR. Both carboxyl groups are formed by the Magnesium-protoporphyrin IX monomethyl ester cyclase, which catalyzes a preceding reaction in the chlorophyll biosynthetic pathway, namely the formation of divinyl-Pchlide. Our computational analysis of the two modes of Pchlide binding reveals different orientations of the crucial carboxyl group at C13^1^. In mode **A**, this group points towards the solvent, while in mode **B**, this group is involved in one of the strongest binding interactions we see between the pigment and the enzyme in both analyzed modes.

Interestingly, previous computational studies have demonstrated that a strong hydrogen bond with the carboxyl group at C13^1^ is responsible for the red shift observed in the fluorescence emission spectrum of the pigment during Pchlide binding to LPOR (12). Our site-directed mutagenesis experiments clearly show that the Y177F mutation allows the enzyme to bind Pchlide, but the resulting complexes exhibit a blue-shifted emission maximum compared to the wild type (Fig. 3BC). Additionally, one of the hydroxyl groups of MGDG is predicted to have a strong interaction with the carboxyl group at C13^1^. Our previous experiments have shown that the addition of MGDG causes a red shift in the emission maximum of the complex (27). These findings suggest that both Y177 and MGDG red shift the emission of the pigment, by forming strong bonds with carboxyl group at C13^1^.

A spectral blue shift in emission, similar to that observed in the Y177F mutant, is also seen in the H319A mutant. However, in the fully assembled complex as determined by cryoEM, the interaction between the residue and the pigment is predicted to be weak and repulsive (Fig. 1D). Interestingly, the H319A mutant is the only mutation discovered thus far that induces a blue shift in the emission maximum of the pigment in the LPOR:Pchlide complex compared to free Pchlide in a buffer solution (Fig. 3C). Previous studies in model systems have demonstrated that the emission maximum of the pigment is influenced by the solvent’s dielectric constant (28). These observations suggest that the binding pocket of the H319A mutant may be more hydrophobic than the water-based buffer. If this is indeed the case, it suggests that in the absence of NADPH, His319 may interact with the pigment, specifically with the carboxyl group at C13^1^. Such an interaction would likely necessitate the enzyme to adopt a different conformation in this complex compared to the structure determined by cryoEM. However, to validate this hypothesis, it is crucial to obtain a detailed structure of the LPOR:Pchlide complex.

An additional interaction specific to mode B, with potential implications for substrate specificity, involves a strong bond between Thr230 and the C13^3^ keto group. This interaction is stabilized by the presence of Lys277, as evidenced by LED analysis (Fig. 1D, 2A). Notably, in a bacterial LPOR variant, mutation of the residue corresponding to Thr230 (*T. elongatus* The147) exhibited a major impact on steady-state activity and NADPH affinity, while only mildly affecting Pchlide binding (29).Supplementary spectral data revealed, however, a blue-shifted absorbance maximum for the mutant complex compared to the WT, indicating a potential variation in the conformation of the binding pocket between the mutant and WT enzyme.

The other identified interactions are not specific to a particular binding mode. For instance, the interaction between the pigment and Gln331, which is most likely a coordination bond, can occur irrespective of the ring’s orientation. Similarly, the hydrophobic interactions involving Phe316, Phe327, and Phe330 are also independent of the binding mode. Mutations of these residues resulted in a significant decrease in pigment binding, confirming their involvement in the process.

Upon comparing the calculated binding energies of mutants with experimental data, we observed a good agreement for Phe237, Phe330, Cys309, and Gln331. However, some noticeable discrepancies were found for Tyr177 and Tyr276. The Y177F mutant exhibited a highly effective Pchlide binding, resulting in no visible band originating for a free pigment in the spectrum (Fig. 3B). Computational predictions, on the other hand, indicate that the bond between Tyr177 and Pchlide contributes to thermodynamically stable pigment binding. It is worth noting that the calculations do not account for the experimental conditions, particularly the presence of lipids, which induce an oligomeric state of the enzyme, causing the pigment to be sequestered in the outer leaflet of the membrane and effectively locked in the complex. The impact of the Y276F mutation is more challenging to explain, as the calculations suggest a weak interaction energy between the residue and Pchlide, implying no effect on Pchlide binding (Fig. 1CD, 2C). However, experimental results demonstrate a significant effect of the Y276F mutation on Pchlide binding (Fig. 3EF). LED analysis uncovered an interaction between Tyr276, Tyr177, and NADPH, suggesting that the observed effect may be attributed to impaired NADPH binding. Nevertheless, our spectroscopic technique is unable to directly detect this interaction.

The findings presented in this paper underscore the significance of two factors that influence the physical properties of Pchlide in complex with LPOR, and should be considered when investigating the enzyme’s reaction mechanism. Firstly, the interactions involving the carboxyl groups of Pchlide have a direct impact on the energy levels of the excited state. Additionally, the orientation of Pchlide within the binding pocket emerges as a crucial determinant. Our results uniformly indicate that the previously assumed binding mode A is disfavored an all counts as compared to mode B, represented in the new cryo-EM structure. The artistic visualization of the process of Pchlide binding in mode B based on pdb:9JK7 is presented in Movie S1. This conclusion appears to contradict the conventional interpretation of the NMR study by Berley and Young, which aimed to identify the carbon atom reduced by NADPH. The atom proposed in that paper, C17 of Pchlide, appears too distant from the pro-S face of NADPH to accommodate direct hydride transfer. Therefore, the mechanism of the light-dependent reaction catalysed by the enzyme therefore remains an open question. A comprehensive analysis of possible reaction pathways incorporating detailed QM/MM simulations at the excited state of the pigment using the structurally-consistent binding mode of the substrate will be necessary. Such analysis should take explicitly into account the critical interactions between the pigment and the LPOR residues highlighted in this work.

## Materials and Methods

### Experiments

LPOR WT and the mutants were expressed in E.coli and purified according to the previously described protocol (27). The AtPORB C309S, AtPORB C309A and AtPORB Y276F mutants have been obtained for previous study (14). The AtPORB F327A_F330A, Q331A and Q331E mutants were obtained using PCR with previously described protocol (27, 30). The primers and temperature conditions are described in supporting information (Table S1).

For spectra measurements, reaction mixtures were prepared according to previously described protocol (27, 31) with protochlorophyllide purified from etiolated wheat seedlings (32). The reaction mixtures were prepared under dim, green light, that was previously shown not to trigger the reaction catalyzed by LPOR, and consisted of: 200 μM NADPH, 5 μM Pchlide, 15 μM LPOR, and the lipids: MGDG:DGDG:PG (50mol%, 35mol%, 15mol%, Avanti Polar Lipids) in a phosphate buffer (37.5 mM phosphate, 225 mM NaCl, 150 mM imidazole, 7 mM 2-mercaptoethanol, 25% w/w glycerol). Samples were incubated for 30 minutes in darkness at room temperature, before being transferred into glass capillaries and frozen in liquid nitrogen. The spectra were measured at 77K with Perkin Elmer LS50b spectrofluorometer set to the following parameters: excitation 440 nm, emission 600-750 nm, slits: 9nm/9nm, 400 nm/min. To verify enzymatic activity, samples were thawed in darkness, illuminated with white light (8 μmol photons m^-2^ s^-1^) for 20 seconds, then frozen in liquid nitrogen, and measured again. For data analysis, the spectra were normalized at maximum intensity. To calculate the photoactive Pchlide as compared to WT, the spectra of free Pchlide, Pchlide in a complex with WT enzyme and the mutants were used. For each, the ratio R between the intensities at 655 nm (emission maximum of oligomeric complex) and at 632 nm (emission maximum of free Pchlide) were calculated. For each experimental condition at least two independent reaction mixtures were analyzed. The photoactive Pchlide was calculated according to the formula: (R_mut_–R_Pch_)/(R_WT_–R_Pch_), where R_x_ is 655/632 intensity ratio of a given mutant (mut), WT enzyme or free Pchlide in a buffer (Pch).

Samples for electron microscopy were negatively stained according to the previously published protocol (14) and visualized with JEOL JEM2100 HT CRYO LaB6 electron microscope. At least two independent samples were analyzed and representative micrographs are presented.

### Computational Methodology

Multiscale simulations were conducted using the MM model, utilizing the cryo-EM structure of LPOR (PDB ID: 7JK9 (14)). A monomer was extracted from the larger oligomeric structure, including the protein and three co-factors (Pchlide, NADP, and LMG lipid) to build the WT and mutants models. To account for missing water molecules in the cryo-EM structure, a systematic hydration protocol was applied, followed by equilibration and 20 ns production simulations. Binding free energy computations were performed using the MM-PBSA protocol (33). QM/MM geometry optimizations were carried out using the ORCA program (34), and LED analysis (21) was conducted on intermolecular interactions in the QM regions in the presence of the protein environment. Additional information on the computational protocols can be found in the supporting information.

## Acknowledgments

We thank Jerzy Kruk for his help with Pchlide purification. The work presented in Fig.2DEF was supported by SONATA project (2019/35/D/NZ1/00295) granted by NCN to MG. This research was funded by the Priority Research Area BioS under the program Excellence Initiative – Research University at the Jagiellonian University in Krakow (B.1.7.2021.16) granted to MG. PP and DAP acknowledge support by the Max Planck Society.

---

**Fig. S1.**
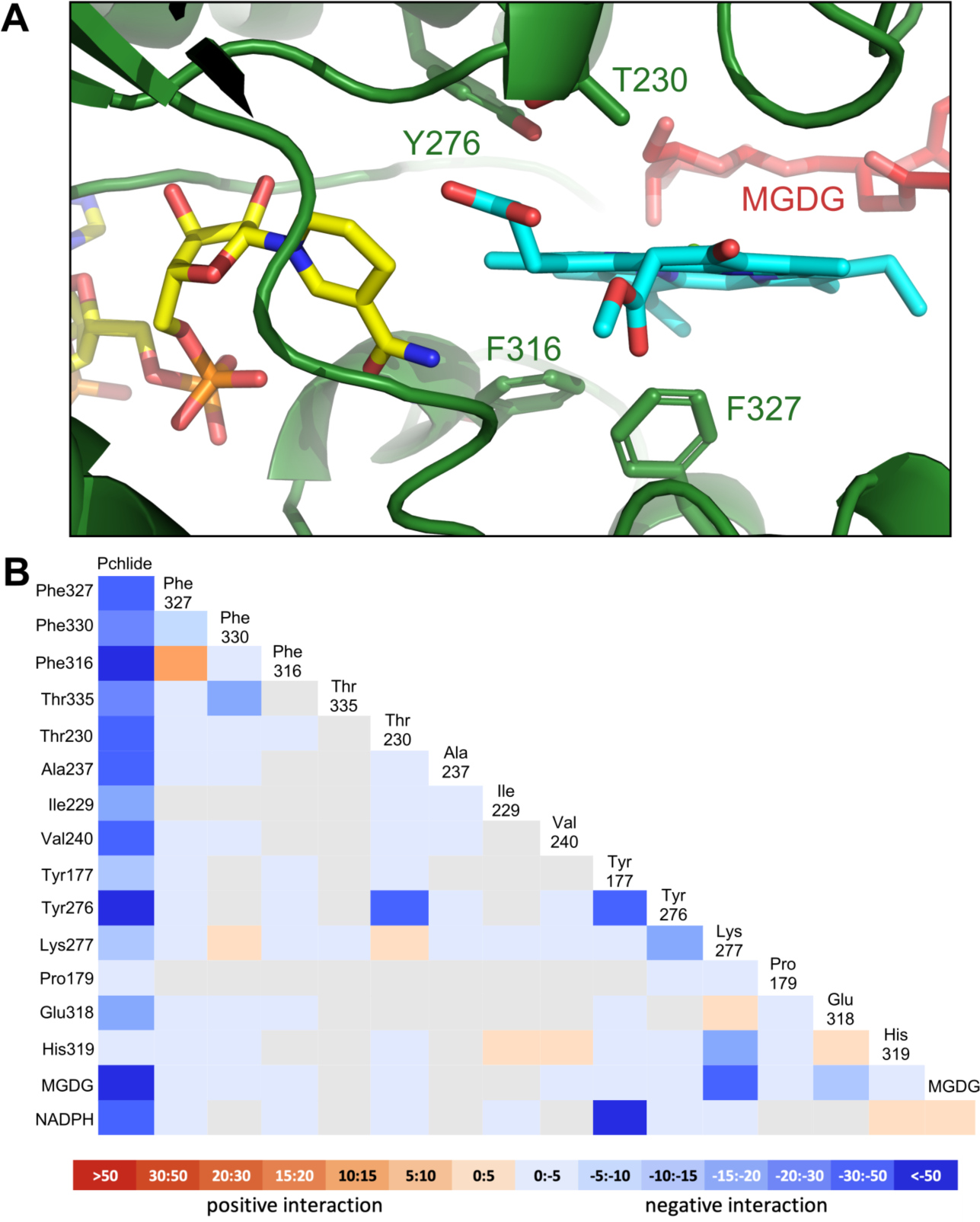
Pchlide binding in mode A. **A**. Detailed view of the pocket with selected residues shown as sticks. **B.** LED interaction map for mode A.

**Fig. S2.**
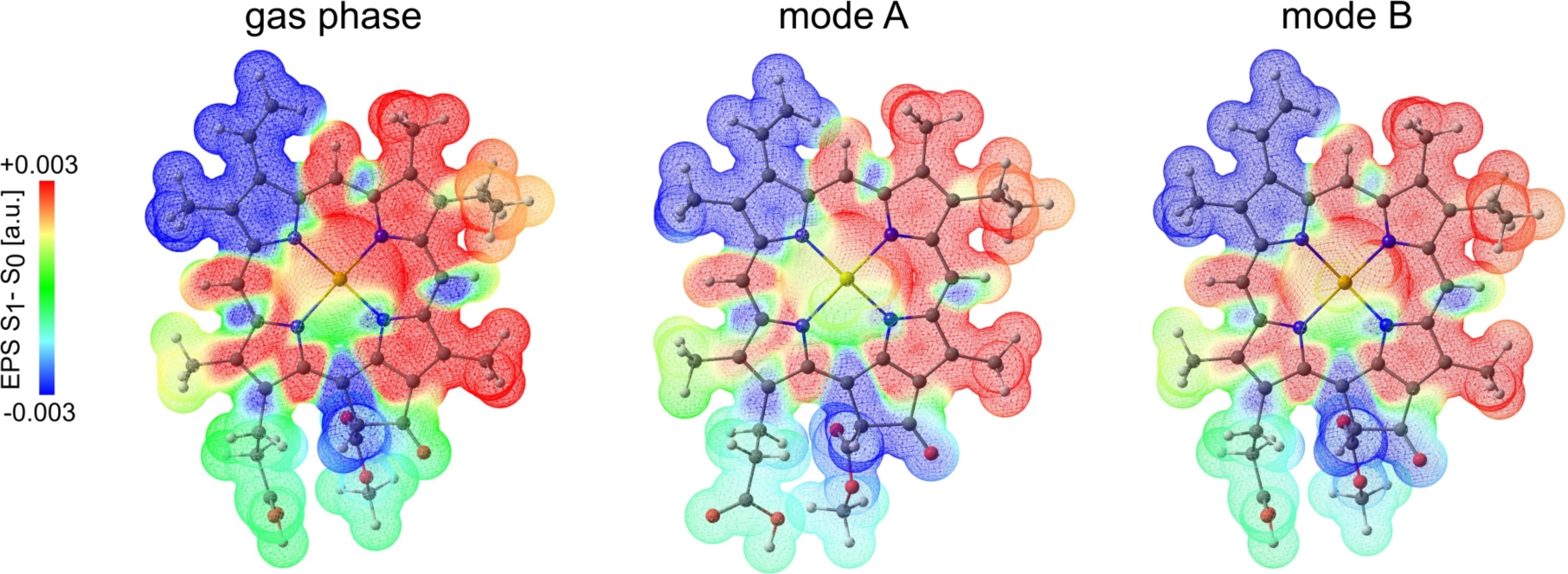
Difference electrostatic potential map (S1 minus S0) associated with the S0 → S1 (Qy) excitation for Pchlide in gas phase, binding mode A and binding mode B.

**Fig. S3.**
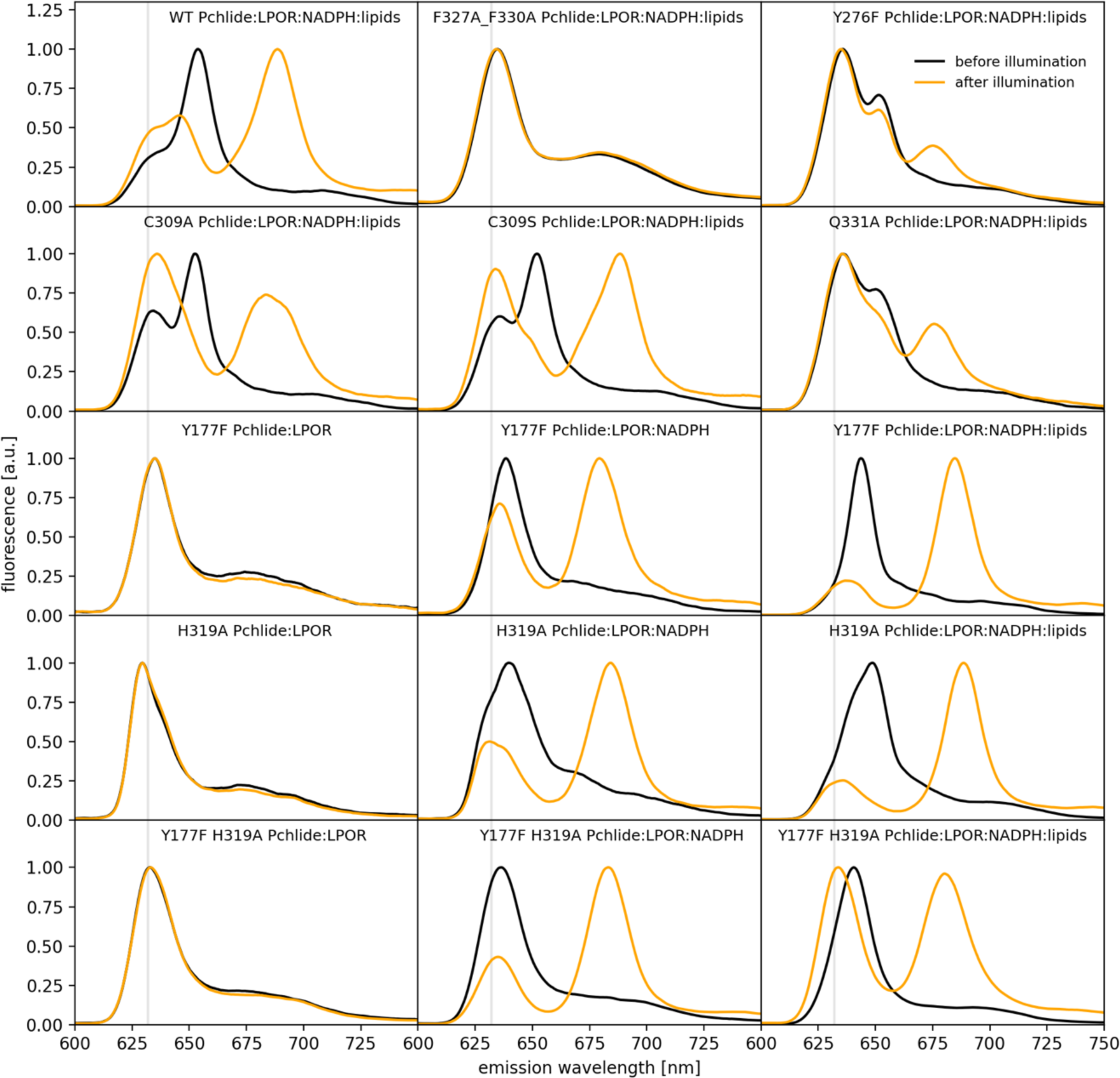
The effect of illumination of the fluorescence spectra of Pchlide in different reaction mixtures. For rows 1 and 2, the reaction was composed of: 15 μM LPOR, 5 μM Pchlide, 200 μM NADPH and 100 μM lipid mix (50mol% MGDG, 35mol% DGDG, 15mol% PG). For rows 3, 4, and 5, specific compositions of the reaction mixtures are provided for each panel; element concentrations mirror those in rows 1 and 2. The vertical grey line denotes the emission maximum of free Pchlide in the buffer.

