## Supplementary information for "Pigment Binding in The Light-Dependent Protochlorophyllide Oxidoreductase"

### Supporting Information for
Pigment Binding in The Light-Dependent Protochlorophyllide Oxidoreductase.

Penelope Pesara, Katarzyna Szafran, Henry C. Nguyen, Abhishek Sirohiwal, Dimitrios A. Pantazis*, Michal Gabruk*

 (M. G.), (D. A. P.)

**This PDF file includes:**

Supporting text
Figures S1 to S3
Tables S1 to S5
Legends for Movies S1
Legends for Datasets S1 to Sx
SI References

**Other supporting materials for this manuscript include the following:**

Movies S1

#### Computational Details

**System Preparation.** The molecular mechanics (MM) model used in this work is based on the cryo-EM structure of LPOR (PDB ID: 7JK9) (1). We extracted a monomer from the large oligomeric structure which included the protein itself and three co-factors (Pchlide, NADPH and MGDG). The water molecules are not resolved, therefore we hydrated the internal cavities using a thorough hydration protocol involving the three-dimensional reference interaction site model (3D-RISM) and integration of the Monte Carlo (MC) method with molecular dynamics (MC/MD) (2-5). Both techniques work in complement to each to other, i.e. 3D-RISM predicts hydration content on static structures which further acts as reference for a dynamic hydration/de-hydration protocol in the MC/MD. This combined strategy has already shown success in predicting the hydration content of membrane-bound Photosystem II (6). The protonation states of protein residues are assigned from the *reduce* program of AmberTools18 (7). The missing tail of MGDG was completed using the *pymol* suite (8). We added two $Na^+$ ions to neutralize the system. After that, the complete system was embedded in a rectangular box and the minimum distance between the solute and box edge was set at 12Å. The entire system contained a total of 72 270 atoms.

The electrostatic charges for the Pchlide, NADPH and LMG co-factors were based on the Merz–Kollman Restrained Electrostatic Potential (RESP) methodology (9). In the first step, we fully optimized the cofactors at the B3LYP/def2-SVP level of theory. After that, single-point HF/6-31G* calculations were performed on the optimized structure and subsequently the RESP fitting of the charges was performed using Multiwfn (4). The bonded parameters for the NADPH and MGDG co-factors were based on the GAFF2 (general Amber force field) force-field (5), while we adapted the Chl *a* force-field for Pchlide parameterization (10). The force-field for the protein was based on the Amber14SB (11), and the TIP3P model (12) was chosen to represent the water. Joung–Cheatham parameters for monovalent ions compatible with the TIP3P water model were employed (13, 14).

**Classical Molecular Dynamics Simulations**. In the first step, all hydrogens atoms were energy-minimized for a total of 2000 steps and all non-hydrogens atoms were restrained to their positions with a force constant of 50 kcal $mol^{-1}$ $Å^{-2}$. In the next step, all atoms of the system were optimized for a total of 2000 steps except the Pchlide, NADPH and MGDG co-factors and backbone atoms (CA, CB, C, O and N) which were restrained with a force constant of 50 kcal $mol^{-1}$ $Å^{-2}$. This restraining force is maintained throughout the simulation protocol. During the equilibration phase, the system is slowly heated from 10 to 300 K within 50 ps, and further propagated for another 50 ps at 300 K in the *NVT* ensemble. The temperature during this procedure is controlled using Langevin dynamics (10) with a collision frequency of 5 $ps^{-1}$. In the next step, the system is propagated for another 1 nanosecond in the *NPT* ensemble. Thereafter, we evoked the MC/MD module implemented in AMBER18. The steric grid for the MC moves was chosen to precisely cover the entire protein volume. The number of Monte-Carlo (MC) move attempts in each MC cycle was set at 1,000,000, whereas the number of MD steps in each MC cycle was set at 1,000. After that, we ran another 10 nanoseconds of simulation in the *NPT* ensemble. During the *NPT* simulations, the collision frequency of the Langevin thermostat is lowered to 1 $ps^{-1}$. The Berendsen barostat (15) was used to regulate the pressure isotropically with a relaxation time of 2 ps and maintained at 1 bar. The Particle Mesh Ewald (PME) approach (16) was used to treat all electrostatic interactions with a 10 Å cut-off. The SHAKE algorithm (17) was used to constrained the bond length involving hydrogen atoms. Integration time-step of 1 fs was used throughout the MD. The energy minimizations were performed using the CPU version while equilibration simulations were performed using the GPU version of the *pmemd* (18-20) engine of the AMBER18 package. The last structural configuration from the MD simulations was taken as input for the QM/MM computations.

**QM/MM Geometry Optimization**. All calculations were performed by employing the additive Quantum Mechanics/Molecular Mechanics (QM/MM) multiscale model with electrostatic embedding as implemented ORCA 5.0. Structural analyses were performed for both binding modes here called mode **A** and **B**. For geometry optimization, density functional theory (DFT) computations were performed using the PBE functional. The size of our system prevents the use of large basis sets, therefore def2-SVP basis set was used for geometry optimizations.
The QM region contains Protochlorophyllide (Pchlide), NADPH, MGDG, and complete amino acids within 6 Å to the central magnesium atom of PChlide. The active MM region (flexible region during QM/MM optimization) contained full residues within a relevant radius of 3 Å to the QM region. The rest of the system was kept fixed during optimization. Due to the extremely high computational cost of DFT in large systems using large QM and active MM regions, the optimization was split up in a stepwise manner. First, NADPH was optimized at the QM level using a charge of -4 for the QM subsystem while the rest of the system was kept fixed. The resulting PDB was taken for the optimization of MGDG at the QM level. Finally, Pchlide and the side chains of important amino acid residues within an approximate distance of 6 Å to the central Mg atom, as well as waters within this radius, were optimized at the QM level with an active region of 3 Å around the QM system including complete residues. This QM region contains full PChlide and the amino acids: Phr327, Phe330, Phe316, Thr335, Thr230, Ala237, Ile229, Val240, Tyr177, Tyr276, Lys277, Pro179, Glu318 and His319 as well as NADPH and MGDG. In order to save computational time, the resolution of the identity approximation was used with the universal auxiliary basis set of Weigend (def2/J) (21).

**Binding Energy Calculations**. The binding free energy corresponding to formation of the LPOR complex, involving Pchlide as the ligand and apo-LPOR as the protein, is determined using the MM-PBSA approach (22-24). This combines Molecular Mechanics, implicit Generalized Born

(GB)/Poisson-Boltzmann (PB) solvation schemes (25) and solvent accessibility surface area calculations to estimate the binding free energy according to equation 1:

$$\Delta G_{bind} = \Delta G_{complex} - (\Delta G_{protein} + \Delta G_{ligand}) \qquad (1)$$

Further theoretical details regarding the MM-PBSA approach can be found elsewhere (23, 26). Here a single-trajectory approach is employed, where only the complex form is propagated, eliminating the need for separate molecular dynamics (MD) simulations for the ligand and protein. All calculations are performed using the parallelized MMPBSA.py module (27) of the Amber/21 package. Solvation free energies are computed using both the Generalized Born (GB) and Poisson-Boltzmann (PB) implicit solvation schemes. Binding energy calculations for the specific system were carried out on a set of equidistant snapshots (total 500), with a temporal interval of 10 ps, extracted from the initial 5 nanoseconds of the NPT simulations. The calculated binding energies were subsequently averaged over this ensemble of snapshots. In order to establish control, we conducted analogous computations over a period of 5–10 nanoseconds (consisting of 500 snapshots), yielding comparable outcomes (see Supporting Information). The solute dielectric constant is set to 2.0 in case of the MM/PBSA computations (28). The entropic contribution ($\Delta S$) to the binding free energy is excluded due to its high computational expense and slow convergence, as the focus is primarily only on the relative binding affinity trends. The calculations are conducted for both orientations of PChlide discussed in the introduction (modes **A** and **B**). Binding energy calculation for the mutants were performed with 'Alanine scanning' functionality of MMPBSA.py module. To gain molecular level insights, binding free-energies were further decomposed into per-residue contributions (29, 30).

**Interaction Energy Calculations and Local Energy Decomposition Analysis**. Local Energy Decomposition (LED) (31-33) analysis at the coupled-cluster DLPNO-CCSD(T) (34-38) was used to analyze the binding in terms of wave function derived interaction energies. This methodology, relying on a local coupled-cluster method with singles, doubles, and perturbatively included triples excitations (34, 37, 39), is a generally applicable interaction decomposition approach (39). Within the LED analysis the system is divided into chemically meaningful fragments, where first the nuclei and subsequently the molecular orbitals (MOs) are assigned to the corresponding fragments. The DLPNO-CCSD(T) interaction energies between Pchlide and a list of surrounding amino acids as well as the lipid MGDG and cofactor NADPH were computed and decomposed according to the LED scheme. Doing so, it is possible to quantify key substrate–residue non-covalent interactions. A similar study has been done by Beck et al. on nicotine and imidacloprid binding to the nicotinic acetylcholine receptor (31).

Within a supramolecular approach, the Enzyme–Substrate (ES) binding energy $\Delta E_{bind}$ can be computed at the DLPNO-CCSD(T) level as the difference between the energy of the ES adduct

($E_{tot}^{SE}$) and that of the isolated E ($E_{tot}^{E}$) and S ($E_{tot}^{E}$) fragments frozen in their in-adduct geometry (31):

$$\Delta E_{bind} = E_{tot}^{SE} - E_{tot}^{S} - E_{tot}^{E} \quad (2)$$

$E_{tot}^{SE}$ can be decomposed into intra-fragment contributions (energy of the substrate $E_{tot}^{E}$ and the energy of the residues $E_{tot}^{R}$ in their adduct geometry) plus a series of inter-fragment contributions, substrate-residue ($E_{tot}^{SR}$) and residue-residue ($E_{tot}^{RR}$) interactions. Similarly $E_{tot}^{E}$ can be decomposed into intra residue $E_{tot}^{E(R)}$ and inter residue-residue $E_{tot}^{E(R,R)}$ contributions. After insertion into (2) and rearrangement, the following equation is obtained describing the binding energy as demonstrated by Beck et al. (31):

$$\Delta E_{bind} = \Delta E_{tot}^{S} + \sum_R \Delta E_{tot}^{R} + \sum_R \Delta E_{tot}^{RR} + \sum_R \Delta E_{tot}^{E(R)} \quad (3)$$

Where $\Delta E_{tot}^{S}$ and $\Delta E_{tot}^{R}$ show the change in energy of the substrate and the residues upon the possible binding modes, and $\Delta E_{tot}^{(R,R)}$ represents the change in residue-residue interaction in the active site.

LED calculations were carried out on the QM/MM models such that all defined fragments used within the LED study were QM-optimized at the PBE/def2-SVP level of theory (35, 37, 38). Single-point DLPNO-CCSD(T) calculations were performed under the electrostatic influence of the protein matrix using def2-TZVP main basis sets with the RIJCOSX approximation and with matching def2/J and def2/C auxiliary basis sets. Due to a relatively large QM region of 348 atoms (link atoms included), VeryTightSCF criterion was applied in combination with LoosePNO settings. The Pipek–Mezey method was used for orbital localization. The DLPNO-CCSD(T) energy was decomposed into a series of additive contributions corresponding to the interaction of pairs of the defined fragments (31-33). The fragmentation includes PChlide as one fragment, as well as NADPH and the lipid MGDG, and a series of amino acid side chains within a distance of 6 Å to the central Mg atom representing other fragments.

**Excited State Calculations and Electrostatic Potential Maps**

Excited state calculations were computed in the framework of time-dependent density functional theory (TD-DFT). Full TD-DFT (without the Tamm–Dancoff approximation) calculations were performed using the hybrid functional B3LYP including 20% Hartree-Fock exchange energy, and range-separated functionals CAM-B3LYP and wB97X-D3BJ which are known to be often superior for spin states and spectroscopic properties. Basis set def2-TZVP and matching auxiliary basis def2/J for employing the RI-J approximation were used for all excited state calculations. Excited state geometries were optimized at the same level of theory. 10 roots were computed and Natural Transition Orbitals (NTOs) were created and visualized for transitions showing significant contributions for the first three excited states using Avogadro. Difference electron density maps were computed for the first three excited states associated with the S0→S1, S0→S2, and S0→S3 transition for all three functionals. The redistribution of electron density upon excitation was studied in gas phase and under the influence of the electrostatic environment of both structural models.

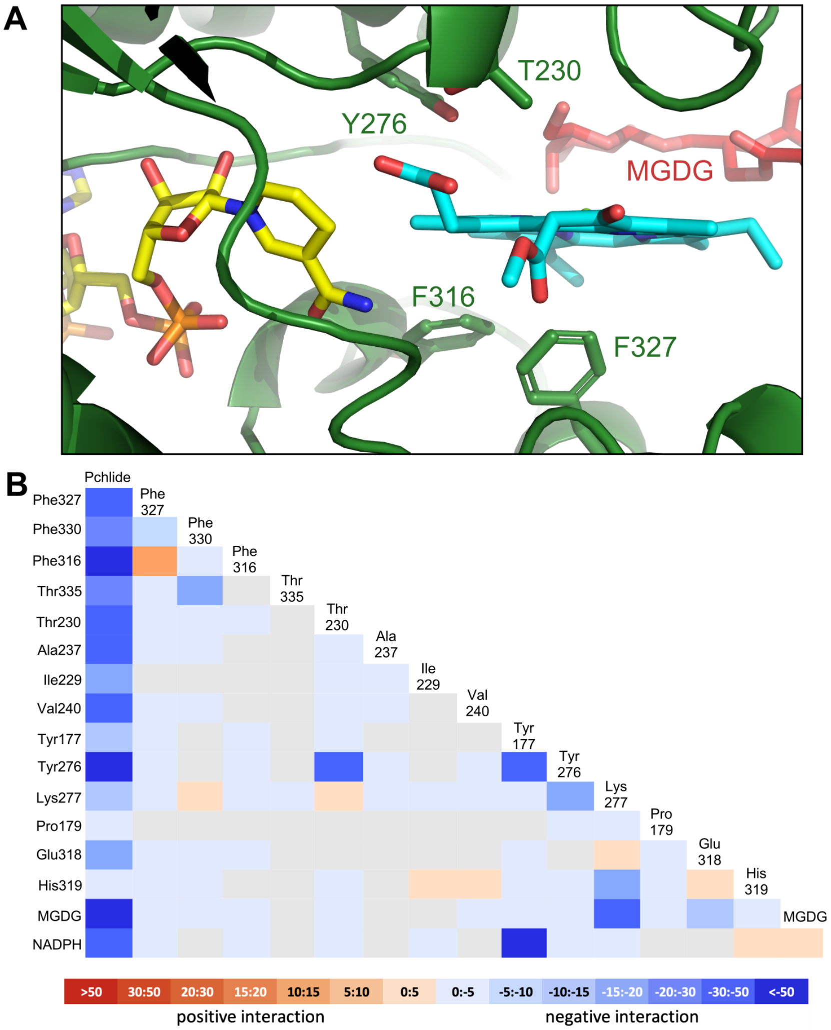


**Fig. S1. Pchlide binding in mode A. A**. Detailed view of the pocket with selected residues shown as sticks. **B.** LED interaction map for mode A.

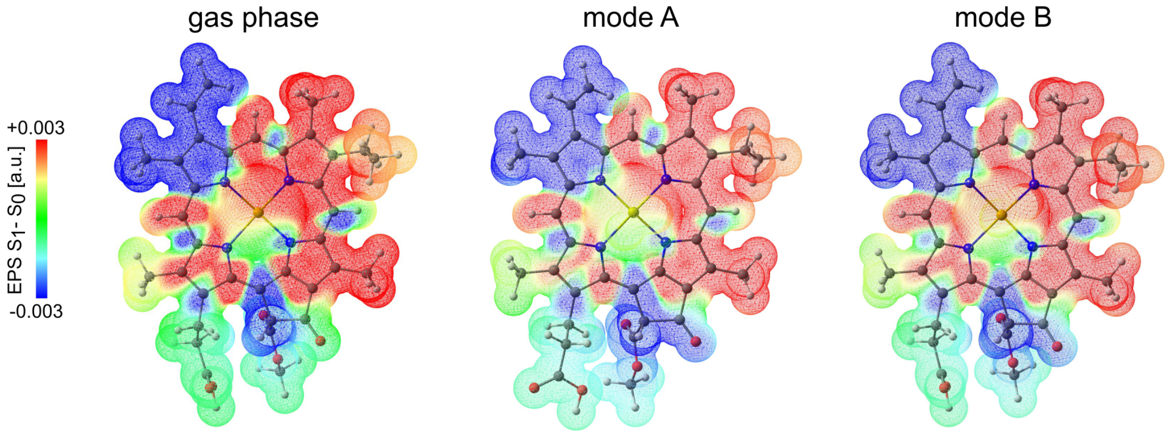


**Fig. S2.** Difference electrostatic potential map ($S_1$ minus $S_0$) associated with the S0 → S1 (Qy) excitation for Pchlide in gas phase, binding mode A and binding mode B.

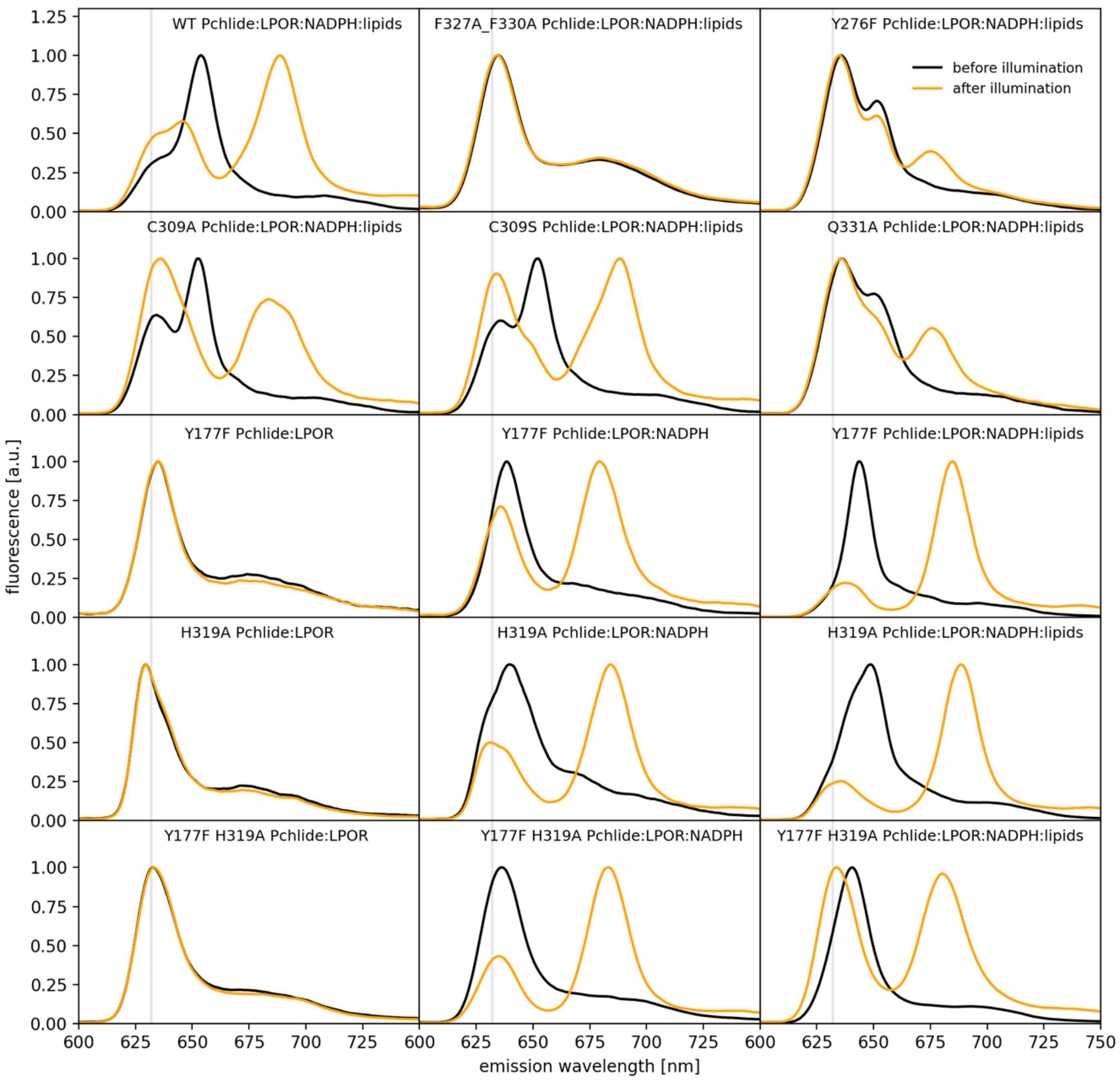


**Fig. S3. The effect of illumination of the fluorescence spectra of Pchlide in different reaction mixtures.** For rows 1 and 2, the reaction was composed of: 15 µM LPOR, 5 µM Pchlide, 200 µM NADPH and 100 µM lipid mix (50mol% MGDG, 35mol% DGDG, 15mol% PG). For rows 3, 4, and 5, specific compositions of the reaction mixtures are provided for each panel; element concentrations mirror those in rows 1 and 2. The vertical grey line denotes the emission maximum of free Pchlide in the buffer.

**Table S1.** List of primers used for site directed mutagenesis.

| Primer | sequence | Annealing temp. [°C] |
|---|---|---|
| F323A_F327A_For_t | GCCCGTGCCCTCGCCCCTCCCTTTCAGAAGTACATC | 63 |
| F323A_F327A_Rev_t | GGCGAGGGCACGGGCGAGAGGAATGTGCTCTCGG | |
| Q331 For | ATCACTAAAGGATATGTCTCCGAAAC | 65 |
| Q331 Rev | GAAGAGGGCACGGAAGAGAG | |
| Q331A_For_t | CCTCCCTTTGCGAAGTACATCACTAAAGGATATGTCT CCG AAAC | |
| Q331A_Rev_t | GTACTTCGCAAAGGGAGGGAAGAGGGCACGGAAGA GAG | |
| Q331E_For_t | CCTCCCTTTGAGAAGTACATCACTAAAGGATATGTCT CCG AAAC | |
| Q331E_Rev_t | GTACTTCTCAAAGGGAGGGAAGAGGGCACGGAAGA GAG | |

**Table S2.** Binding energy of PChlide computed using the MM-PBSA and MM-GBSA approaches for both binding orientations of the pigment. Energy decomposition of the binding energy (i.e., Complex - Receptor - Ligand) is also provided.

| | | vdW | EEL | $\Delta G_{gas}$ | $\Delta G_{solv}$ | $\Delta G_{bind}$ |
|---|---|---|---|---|---|---|
| **PB** | **Mode B (Cryo-EM)** | -61.4855 | -7.3492 | -68.8347 | 66.3703 | -2.4644 |
| | **Mode A** | -47.4520 | -9.2073 | -56.6592 | 62.9498 | +6.2906 |
| **GB** | **Mode B (Cryo-EM)** | -61.4855 | -14.6984 | -76.1839 | 30.8558 | -45.3281 |
| | **Mode A** | -47.4520 | -18.4145 | -65.8665 | 38.6871 | -27.1794 |

**Table S3.** Binding energy (in kcal/mol) of PChlide computed using the MM-PBSA and MM-GBSA approaches for both binding orientations of the pigment. The calculations were performed using snapshots from 0–5 ns and 5–10 ns part of the MD trajectories. A solute dielectric constant of 2 is used in MM-PBSA.

| Approach | Trajectory | Mode B | Mode A |
|---|---|---|---|
| **MM-PBSA** | **0–5 ns** | -2.4644 | 6.2906 |
| | **5–10 ns** | -2.8591 | 6.3334 |
| **MM-GBSA** | **0–5 ns** | -45.3281 | -27.1794 |
| | **5–10 ns** | -44.8850 | -27.3387 |

**Table S4.** Dependence of the solute dielectric constant on the MM-PBSA binding energy (i.e., Complex–Receptor–Ligand, in kcal/mol). All computations were performed for the snapshots derived from the 0–5 ns part of the full trajectory.

| Dielectric constant | Trajectory | Mode B (Cryo-EM) | Mode A |
|---|---|---|---|
| **1** | **0-5 ns** | 21.5171 | 27.0882 |
| **2** | **0-5 ns** | -2.4644 | 6.2906 |
| **4** | **0-5 ns** | -14.3125 | -4.0543 |

**Table S5.** Total free-energy decomposition into per-residue contributions for mode **A** and mode **B**. Only residues with a significant interaction to the substrate PChlide in one of the two binding modes over/under a defined threshold (*x*>0.1; -0.1>*x*) are shown. All energies in kcal/mol.

| Residue | MODE A | MODE B |
|---|---|---|
| Tyr 177 | -0.55 | -3.55 |
| Pro 179 | -0.45 | -0.31 |
| Ser 228 | 0.52 | -0.21 |
| Ile 229 | 0.49 | -0.83 |
| Thr 230 | -2.20 | -2.96 |
| Gly 231 | 0.16 | 0.01 |
| Asn 232 | 0.12 | 0.05 |
| Thr 235 | -0.09 | -0.12 |
| Leu 236 | -0.25 | -1.18 |
| Ala 237 | -1.20 | -2.01 |
| Val 240 | -1.51 | -1.53 |
| Pro 242 | -0.21 | -0.27 |
| Lys 243 | 0.18 | 0.20 |
| Asp 271 | -0.17 | -0.27 |
| Ala 273 | -0.05 | -0.60 |
| Lys 274 | 0.12 | 0.06 |
| Tyr 276 | -0.94 | -1.17 |
| Lys 277 | 0.62 | 0.46 |
| Asp 278 | -0.11 | -0.16 |
| Lys 280 | 0.95 | 0.95 |
| Tyr 306 | -1.17 | -0.08 |
| Gly 308 | -0.36 | -0.30 |
| Cys 309 | -0.82 | -1.27 |
| Ile 310 | -0.12 | -0.10 |
| Leu 315 | -0.35 | -0.91 |
| Phe 316 | -1.58 | -0.91 |
| His 319 | -0.12 | -0.09 |
| Phe 323 | -0.62 | -0.82 |
| Phe 326 | -0.12 | -0.12 |
| Phe 327 | -2.34 | -2.14 |
| Phe 330 | -0.89 | -1.11 |
| Gln 331 | -1.28 | -1.83 |
| Thr 335 | -0.43 | -1.60 |
| Lys 336 | 0.16 | 0.16 |
| Gly 337 | -0.21 | -0.44 |
| Lys 338 | -0.11 | -0.05 |
| Arg 347 | 0.12 | 0.11 |
| Tyr 363 | -0.20 | -0.01 |
| Asn 375 | 0.10 | 0.04 |
| NDP 403 | 2.45 | 1.59 |
| LMG 404 | 0.35 | 0.19 |

**Movie S1.** The artistic visualization of the process of Pchlide binding in mode B based on pdb:9JK7.
